# Resident memory T cells in dirty mice suppress innate cell activation and infiltration into the skin following stimulation with alarmins

**DOI:** 10.1101/2024.07.11.602963

**Authors:** Meaghan E. Story, Laura K. Ferris, Alicia R. Mathers

## Abstract

Trm cells are sequestered at barrier tissues as a swift first line defense against peripheral reinfections in both antigen dependent and antigen independent bystander modes. Trm cells are also capable of mediating autoimmune diseases, such as psoriasis, wherein autoreactive Trm cells are aberrantly activated. To quickly combat infections, activated Trm cells can stimulate the influx and activation of memory T cells and innate immune cells. However, there is significant heterogeneity in the inflammatory responses that Trm cell populations can induce, specifically in the activation of the innate profile. Most studies to date have utilized a reductionist approach to examine single Trm populations, specific pathogens, and defined tissues. Herein, we adopted a more holistic approach utilizing barrier-free ‘dirty’ mice to profile activated innate cells attracted to the skin in the presence of quiescent cutaneous Trm cells. Notably, dirty mice are a more human predictive model due to having a diverse microbial experience that leads to the development of a complete complement of Trm cells in the skin. We demonstrate that in the dirty mouse model mice have a significant reduction in cutaneous neutrophils and monocytes compared to SPF mice following local treatment with two separate innate stimuli. These findings reveal that cutaneous Trm cells have the capacity to temper the innate immune response and further substantiate the implication that Trm cells are heterogenous in their functions depending in large part on their tissue residency. However, in an autoimmune microenvironment Trm cells are capable of recruiting innate cells to the site of an exposure to a damage-associated molecular pattern. Likely due to the imbalance of IL-17 and IFN-γ.

## Background

Trm cells are sequestered at barrier tissues as a swift first line defense against peripheral reinfections by secreting pro-inflammatory and cytotoxic mediators.^1-3^ In addition to their antimicrobial properties, Trm cells are also capable of mediating autoimmune diseases, such as psoriasis, wherein autoreactive Trm cells are aberrantly activated.^4,5^ As an additional mechanism to quickly combat infections, activated Trm cells can stimulate the influx and activation of central memory T cells (Tcm) and innate immune cells.^1,2,6,7^ However, while Trm cells from different tissues have a similar transcriptome there appears to be more heterogeneity in the Trm cell populations than originally believed.^4,8,9^ In this regard, in specific pathogen free (SPF) mice following viral rechallenge CD8^+^ Trm cells in the female reproductive tract (FRT) and oral mucosa, induce dendritic cell (DC) maturation and attract NK cells.^1,10^ In the lungs, a pulmonary infection with bacteria led to the bystander activation of CD8^+^ Trm cells, which triggered a significant increase of neutrophil infiltration into the airways.^2^ Whereas, in skin populated by CD8^+^ Trm cells, Ariotti et al.^3^ observed an increase in neutrophil infiltration only upon specific peptide rechallenge and not by bystander activation. Glennie et al. ^11^ demonstrated that CD4^+^ Trm cells mediate inflammation following a cutaneous *Leishmania major* rechallenge by recruiting inflammatory monocytes and not neutrophils or DCs into the skin. Hence, the effects Trm cells have on the innate immune response in the respective tissues are still poorly defined.

Regardless of whether the Trm cells are activated by cognate antigen in a canonical mode or by bystander activation in a non-canonical mode, both Trm cell populations produce IFN-γ and can exert cytotoxicity upon activation.^2,3,6^ IFN-γ-producing Trm cells can maintain IFN-γ expression long after inflammatory resolution when Trm cells have entered a quiescent state.^1-3,7^ Following antigen rechallenge or bystander activation IFN-γ levels are then significantly increased to levels capable of promoting effector functions. Shenkel et al.^6^ demonstrated that Trm cells recruit central memory T (Tcm) cells into the reproductive track in an IFN-γ-dependent manner. In this case, enhanced IFN-γ levels stimulated cells in the microenvironment to express chemokines such as CXCL9 (MIG) and CXCL10 (IP-10), which are lymphocyte chemoattractants. Moreover, following cutaneous rechallenge with *L. major*, Trm cells produce increased levels of IFN-γ and chemoattract circulating Tcm cells in a CXCR3-dependent manner.^7^ Intriguingly however, IFN-γ also suppresses innate myeloid cell chemotaxis by blocking their ability to respond to CCL2, CCL3, and CCL4 in an apparent effort to limit tissue inflammation.^12^ Likewise, type I Interferon and IFN-γ block neutrophil infiltration into ganglia by suppressing CXCL2.^13^ However, the levels of IFN-γ expressed by different Trm cell populations can be variable. In this regard, cutaneous CD8^+^ Trm cells expressing CD49a and CD103 produce IFN-γ, perforin, and granzyme B upon activation and exposure to IL-15, whereas activated CD8^+^ Trm cells expressing only CD103 produce IL-17 and IFN-γ to a lesser extent.^4^ Thus, further studies are necessary to gain a deeper understanding of how Trm cells direct the innate immune response. Importantly, studies are lacking that address the effects quiescent Trm cells have on the innate response following murine exposure to sterile innate stimuli.

Here we utilized barrier-free ‘dirty’ mice (DM) to assess the capacity of quiescent IFN-γ producing cutaneous Trm cells to attract innate cells into the skin. Notably, DM are a more human predictive model due to having a diverse microbial experience that leads to the development of a complete complement of Trm cells in the skin.^4,14,15^ We demonstrate that in the DM model mice have a significant reduction in cutaneous neutrophils and monocytes compared to SPF mice following local treatment with two separate innate stimuli. These findings reveal that cutaneous Trm cells have the capacity to temper the innate immune response and further substantiate the implication that Trm cells are heterogenous in their functions depending in large part on their tissue residency. However, in an autoimmune microenvironment Trm cells are capable of recruiting innate cells to the site of an exposure to a damage-associated molecular pattern (DAMP). Likely due to the imbalance of IL-17 and IFN-γ.

## Methods

### Animals

For DM studies, female mice were acquired from a local pet store in Pittsburgh, PA and housed in disposable caging in a BSL-3 facility. Mice were then housed under specific-pathogen free conditions. All mice were treated according to the University of Pittsburgh’s Institutional Animal Care Guidelines and the NIH guide for the care and use of laboratory animals. To induce innate activation, mouse backs were shaved or treated with a depilatory cream then injected intradermally (i.d.) with 100 μl of BzATP (350 μmol/L) (Sigma-Aldrich, St. Louis, MO) in combination with POM1 (3.2 mg/kg) (Tocris Bioscience, Bristol, UK), in PBS (vehicle control) at two sites daily for 5 consecutive days then every other day (EOD) for 4 days and sacrificed on day 10. In separate experiments, innate inflammation was induced in mice with 62.5 mg of IMQ (5% Aldara; 3M Pharmaceuticals, St. Paul, MN) topically administered on the backs and ears daily for 5 days. For xenotransplants, NOD-*scid* IL2Rgamma^null^ (NSG) mice (The Jackson Laboratory; # 005557) were used. The NSG mice were treated with 150 μg (i.p.) mouse Gr-1 specific antibody (Bio X Cell; Lebanon, NH) on day -1 of transplant and then every 4-5 days after transplant to neutralize murine granulocytes^16^.

### Human Skin xenotransplants

Human skin biopsies were used for xenotransplants. For this, non-lesional psoriasis skin shave biopsies were obtained through the UPMC Dermatology clinic from psoriasis patients with moderate to severe plaque-type psoriasis that were treatment naïve. All human samples were procured with informed consent in accordance with the Declaration of Helsinki protocols and University of Pittsburgh Institutional Review Board approval. The resulting human skin samples were comprised of the epidermis and a thin layer of underlying dermis with a thickness of approximately 0.05-1.0 mm. Shave biopsies were cut into 10-15 mm diameter grafts and orthotopically grafted onto the backs of anesthetized NSG mice. Grafts were allowed 2 wks for acceptance before injections with indicated treatments.

### Real-time quantitative RT-PCR

Total RNA was extracted from tissues utilizing TRI-reagent (Molecular Research Center, Cincinnati, OH), quantified utilizing a NanoDrop (DeNovix, Wilmington, DE), and reverse-transcribed using the QuantiTect Reverse Transcription Kit according to manufacturer’s instructions (Qiagen, Hilden, Germany). Real-time quantitative PCR was performed utilizing Taqman Gene Expression Master Mix (Life Technologies, Carlsbad, CA) according to manufacturer’s instructions. Taqman primers utilized were specific for Rpl37a (endogenous control), IL-1β, IL-6, and IL-23. Reactions were run in triplicates and analyzed on a QuantStudio^™^ 6 Pro sequence detection system (Applied Biosystems, Waltham, MA) equipped with a 384-well block. Relative fold-changes of RNA transcript expression levels were normalized based on the 2^-ΔΔCt^ method.

### Tissue Cytokines

At indicated time points, skin samples (4 mm) were collected, minced, and placed into Cell Lysate Buffer (RayBiotech, Norcross, GA) supplemented with protease inhibitors. Lysates were diluted 1:2, and cytokine concentrations were measured in duplicate using Luminex technology with the Fluorokine Multianalyte Profiling kit according to manufacturer’s instructions (R&D Systems, Minneapolis, MN). Samples were read on a Bio-Plex 200 system (BioRad, Hercules, CA) using the Bioplex 6.1 software.

### Spectral Flow Cytometry

Back skin biopsies (10 mm) were collected, minced, and enzymatically digested in 1 mg/ml Collagenase D (Roche, Indianapolis, IN), 1 mg/ml DNAse (Roche), 10 mg/ml hyaluronidase (Sigma-Aldrich), and 0.1% BSA in IMDM (ThermoFisher) for 45 minutes at 37 °C. Next, 10 mmol/L EDTA was added for an additional 5 minutes at room temperature. To make a single-celled suspension, samples were passed over a cell strainer (Corning, Corning, NY). The following antibodies were purchased from BioLegend (San Diego, CA): BV421 CD103 (clone QA17A24), BV510 or PerCP/Cy5.5 CD45 (clone 30-F11), BV711 CD3ε (clone 145-2C11), BV785 CD69 (clone H1.2F3), PE CD4 (clone GK1.5), Alexa Fluor 647 CD8a (clone 53-6.7), BV785 CD11b (clone M1/70), Alexa Fluor 488 Ly6C (clone HK1.4), P2/Cy7 CD11c (clone N418), Alexa Fluor 647 Ly6G (clone 1A8). The fixable viability dye eFluor 780 was purchased from eBioscience (San Diego, CA). To block non-specific binding through Fc receptors anti-CD16/CD32 (clone 2.4G2; BD biosciences) was used. Cells were analyzed using a Cytek® Aurora spectral flow cytometer (Bethesda, MD) and FlowJo software (BD; Ashland, OR) was utilized for analysis. The gating strategy was to gate on CD45 positive cells versus viability dye negative cells, doublet exclusion, and then on living cells in FSC versus SSC.

### Microscopy

Cross-sections of mouse ears and back were prepared and stained as previously described.^17^ Briefly, cross-sections were embedded in Tissue-Tek OCT (Miles Laboratories; Elkhart, IN), snap frozen in pre-chilled methyl-butane (Sigma-Aldrich), and stored at − 80 °C until ready to use. Cryostat sections (8 µm) were mounted onto slides pre-treated with Vectabond (Vector Laboratories; Burlingame, CA), air-dried, and fixed in cold 96% EtOH (10 min) and used for H&E staining or immunofluorescence labeling. For immunofluorescence, tissue sections were blocked with 10% normal goat or donkey serum in PBS and the avidin/biotin blocking kit (Vector Laboratories) if necessary, and immunofluorescently labeled with combinations of the following specific antibodies: Alexa Fluor 488 CD11b (Biolegend; clone M1/70), Alexa Fluor 594 Ly6G (Biolegend; clone 1A8),

Alexa Fluor 647 LY6C (AbD Serotec; Raleigh, NC; clone ER-MP20). Nuclei were counterstained with DAPI (Molecular Probes, Eugene, OR). Images were acquired using a Keyence BZ-X800 fluorescence microscope (Itasca, IL) and quantitated with Keyence Hybrid cell count software module, which can determine colocalization of fluorescent signal.

### Statistics

All results were analyzed with GraphPad Prism10 Software (San Diego, CA). Results from multiple different groups were compared using a one-way analysis of variance (ANOVA) followed by Tukey’s multiple comparison post-hoc test. Comparison of two means was performed by a 2-tailed Student’s T-test. A *p* value < 0.05 was considered statistically significant. \**p* < 0.05, \*\**p* < 0.01, \*\*\**p* < 0.001, \*\*\*\**p* < 0.0001.

## Results and Discussion

To assess the role that Trm cells have on directing innate immunity in the skin, we have utilized pet store mice, a DM model of Trm cells.^14,15^ The DM model has a Trm compartment that consists of a vast number of Ag-experienced Trm cells that more closely models adult human immune responses, making it an excellent holistic model to study Trm cells *in vivo*.^14,15^ However, it was necessary to first confirm the increase of cutaneous CD4^+^ and CD8^+^ Trm cells in our DM cohorts. In this regard, we stained cutaneous single-cell suspensions from naive SPF mice and DM for CD4^+^CD69^+^CD103^+^ and CD8^+^CD69^+^CD103^+^ Trm cells and observed a significant increase in both CD4^+^CD69^+^ and CD8^+^CD69^+^ Trm cells in DM cohorts, compared to SPF mice (**Figure 1**). Additionally, based on third color staining we also detect CD103^int/high^ and CD103^int^ populations of Trm cells (**Figure 1**). Consistent with previous findings, cutaneous Trm cells have variable expression of CD103, a molecule important for epidermal retention.^4^ Interestingly, it has been suggested that CD8^+^CD103-Trm cells in the epidermis become activated upon challenge and migrate into the dermis.^4^ Thus, DM have an expansive cohort of Trm cells in the skin capable of maintaining cutaneous Trm residency and responding to potential threats. Moreover, these findings are consistent with the populations and ratios of CD4 and CD8 Trm populations identified in normal human skin.^18^.

**Figure 1.**
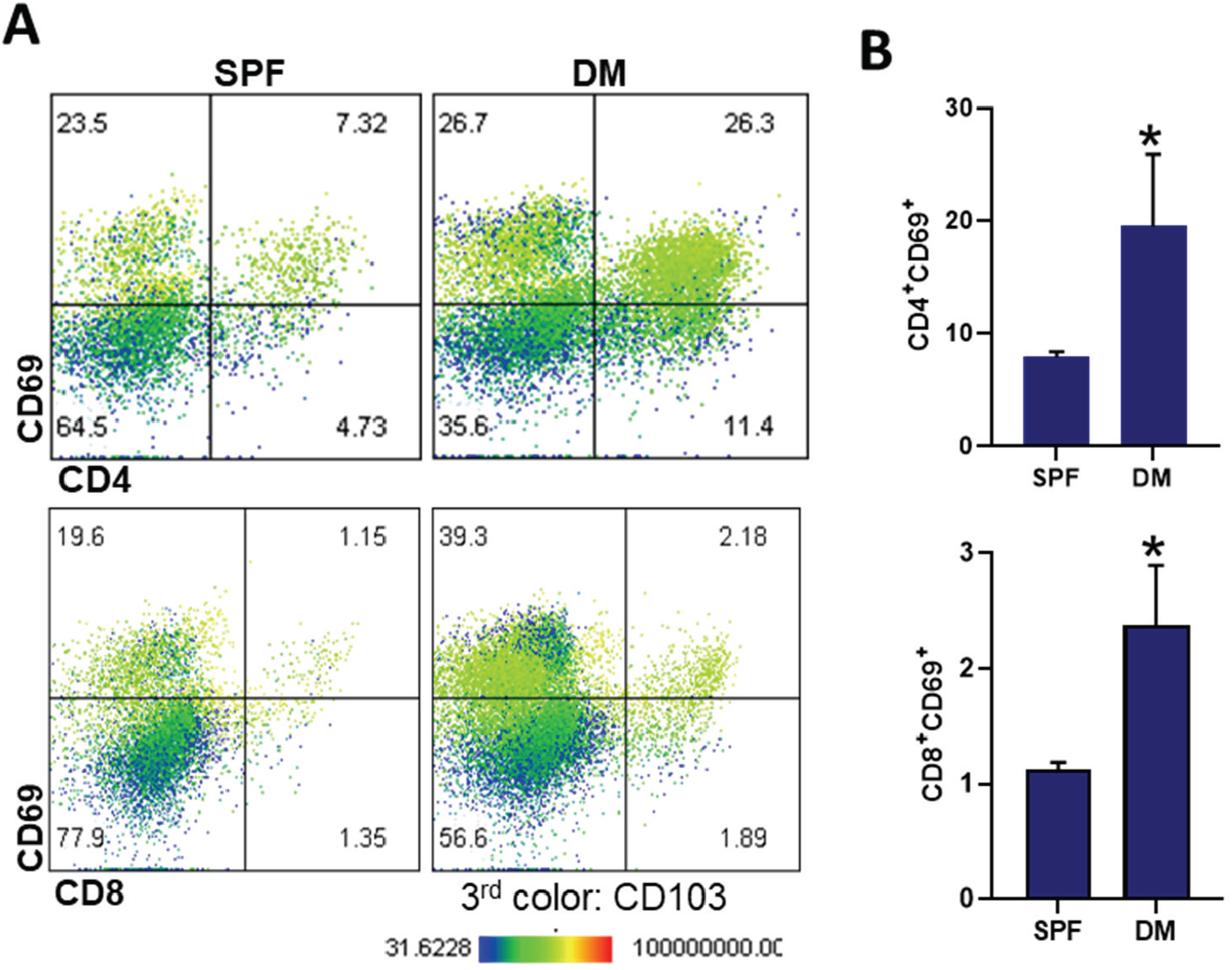
DM have a significant increase in both CD4 Trm and CD8 Trm cells in the skin. Mouse skin was collected from untreated pet store (PS) and SPF (C57Bl/6) mice and single-cell suspensions from each individual mouse was stained and analyzed by Aurora Flow cytometry. **(A)** Representative dot plots displaying analysis of CD4 and CD8 versus CD69 and 3 -color CD103 analysis. Cells were initially gated on CD45 Via, doublet exclusion, and FSC vs SSC. **(B)** Quantitation of the Flow cytometry data. Bars are the mean ± SEM from 3-5 individual mice (DM; n=5 and SPF n=3).

To evaluate the capacity of DM to mount an innate immune response, we utilized the imiquimod (IMQ) model of cutaneous inflammation. IMQ is a pathogen-associated molecular pattern (PAMP) and TLR 7/8 ligand capable of inducing cutaneous inflammatory responses when applied topically.^19,20^ SPF mice and DM received a topical dose of IMQ cream (5% Aldara) or control cream (Vanicream, VC) on the shaved backs and ears for 6 consecutive days.^19,21,22^ On day 7, cross-sections from ear (**Figure 2A and 2B**) and back skin (**Figure 2C**) were collected. A histological assessment determined that there was a significant increase in epidermal thickness in SPF mice and DM, compared to VC treated controls (**Figure 2C**). However, when comparing DM to SPF mice, there was a significant decrease in epidermal thickness in DM treated with IMQ, compared to SPF mice (**Figure 2C**). A quantitative analysis of the cutaneous infiltrate revealed a decrease in CD11b^+^ cells in DM, compared to SPF mice (**Figure 2B**). Notably, there was a significant decrease in CD11b^+^Ly6G^+^ neutrophils and CD11b^+^Ly6C^+^ inflammatory monocytes in DM, compared to SPF mice (**Figure 2B**). Fluorescent intensity of antibody staining was also assessed and was consistent with cell counts (data not shown). Finally, a transcript assessment revealed a significant increase in the innate inflammatory cytokines, IL-β, IL-6, and IL-23 in SPF mice compared to DM (**Figure 3)**. Thus, there was a significant decrease in the collective innate inflammatory response induced in DM compared to SPF mice.

**Figure 2.**
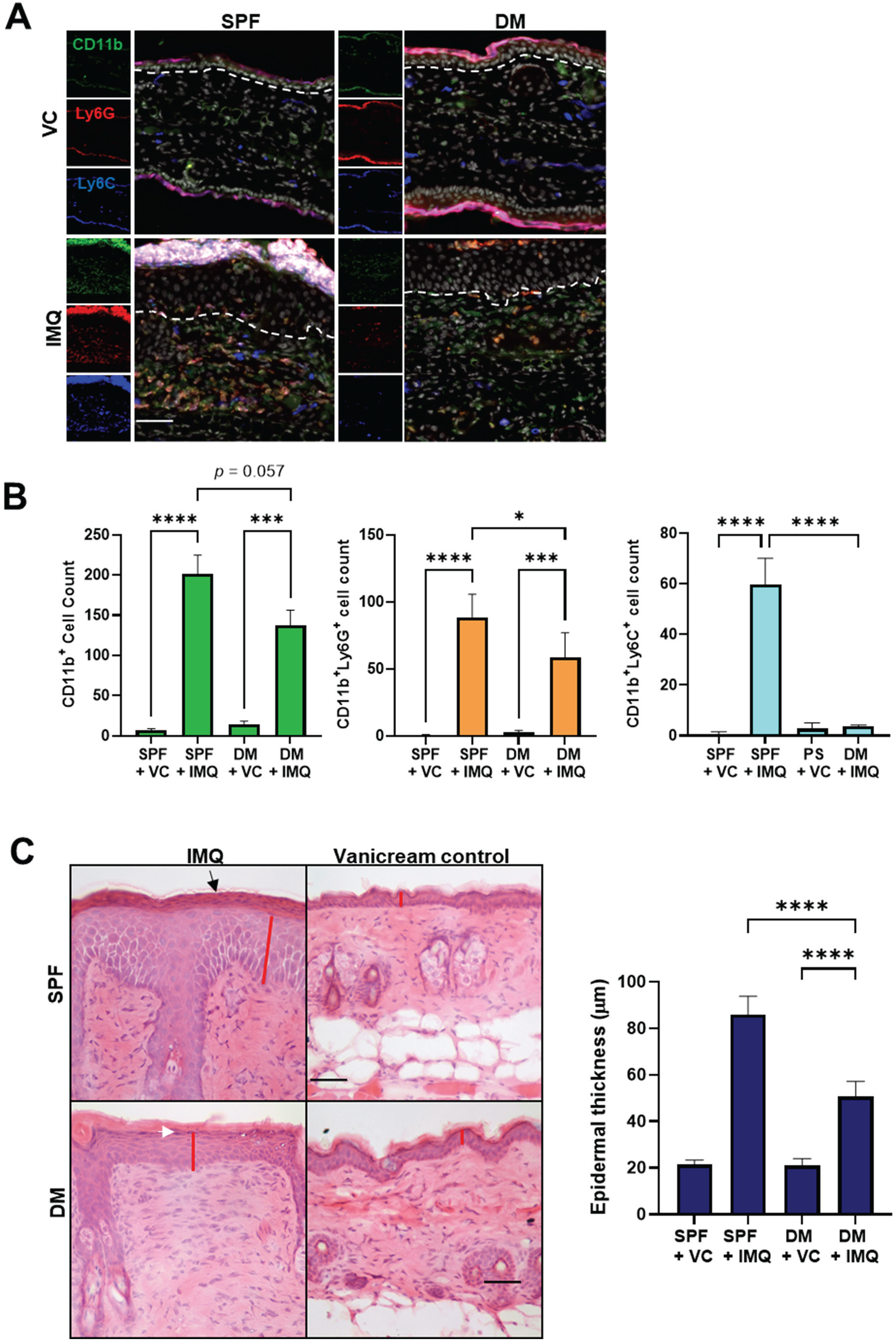
In dirty mice, cutaneous inflammation is decreased following treatment with IMQ. SPF and DM were treated on the ears and back with IMQ or Vanicream, vehicle control (VC), for 6 days. On day 7 cutaneous cross-sections were collected and (**A**) immunofluorescently labeled with CD11b-(green), Ly6G-(red), and Ly6C-specific antibodies (blue) in addition to DAPI nuclei counterstain (white). Dashed line indicates the epidermal-dermal junction. (**B**) Bars represent the mean ± SEM (n=3-4) of the Total CD11b cells, CD11b Ly6G neutrophils, and CD11b Ly6C monocytes. 5-8 sequential non-overlapping sections were averaged from each mouse. (**C**) H&E cross-sections from back skin. Black arrow indicates parakeratosis (cell nuclei retained in the stratum corneum keratinocytes), white arrow indicates the maintenance of the granular layer, and the red line indicates epidermal thickness. **(D)** Bars represent the mean ± SEM (n=4) of epidermal thickness from. 10 measurements were averaged from each mouse. (**D**) Bars represent the mean ± SEM (n=4) of Fold-change of IL-1β transcripts. Each sample was run in triplicate.

**Figure 3.**
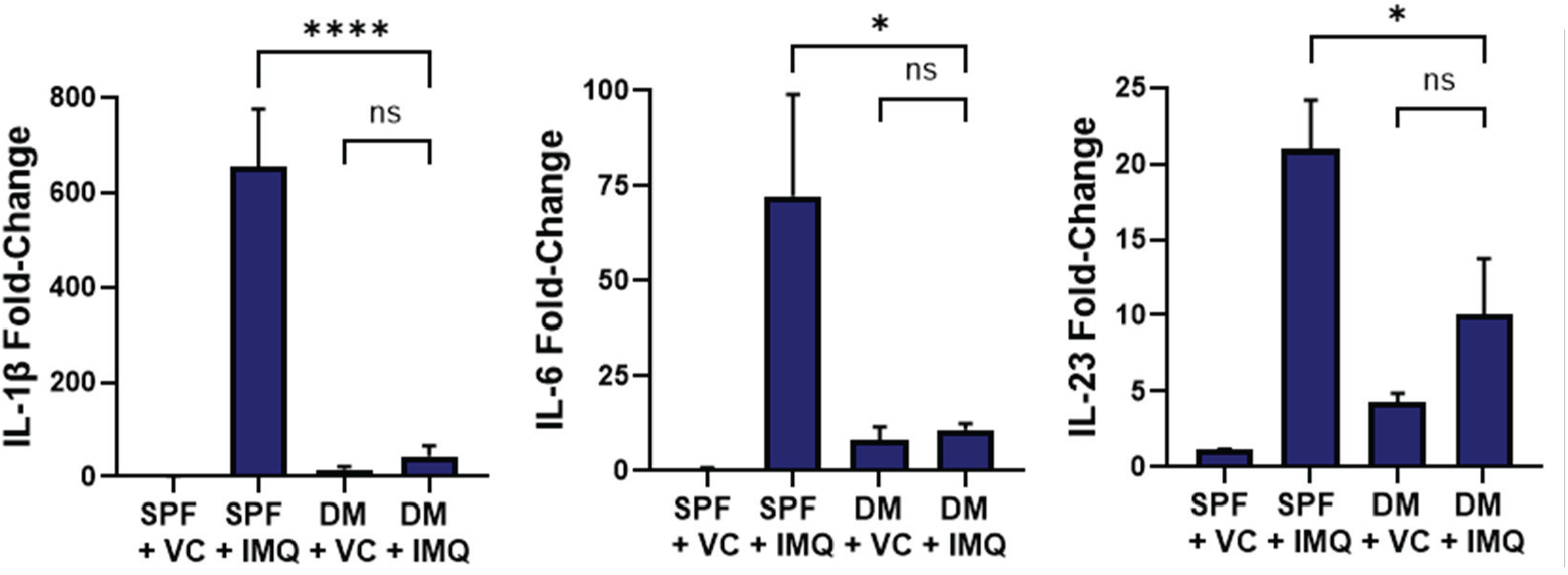
Innate cytokine transcripts are significantly decreased in DM. SPF and DM were treated on the backs with IMQ or Vanicream, vehicle control (VC), for 6 days. On day 7 back skin was collected and assessed for IL-1β, IL-6, and IL-23 innate cytokine transcripts. Bars represent the mean ± SEM (n=4) of indicated transcripts. Fold-change was determined using qRT-PCR and the 2^-ΔΔCt^ method. Each sample was run in triplicate.

Interestingly, our results are not in line with prior publications. There are likely many factors that may contribute to these differences. In the current studies, we utilized a more holistic approach to mimic the human response to a greater extent. However, prior studies used a more reductionist approach to isolate 1) the role of either CD4^+^ Trm cells or CD8^+^ Trm cells, 2) individual pathogens, 3) antigen-specific versus bystander activation, 4) viral, bacterial, or parasite challenge, or 5) tissue residency. However, while both approaches provide useful insight into the effects Trm cells have on the innate immune response, it is abundantly clear that more studies are needed to reliably integrate findings from both approaches.

Repeated exposure to toll-like receptor (TLR) agonists can result in receptor desensitization and induction of immune tolerance.^23,24^ Thus, to rule out a role for TLR-tolerance induced by the microbiome and to confirm our prior results, we subsequently employed an ATP DAMP model that is a non-tolerizable model. ATP signals through the purinergic P2X7 receptor (P2X7R), a purinergic receptor and ion channel that when triggered leads to large pore formation, Ca^2+^ influx, inflammasome assembly, translocation of NF-κB, and induction of inflammatory cytokines. Specifically in the skin, the P2X7R-induced inflammatory response is characterized by a significant increase in the inflammatory mediators IL-1β, IL-1α, TNFα, IL-6, IL-23, and IL-17 and a significant increase in DC, monocyte, and neutrophil infiltration.^21^ Thus, the activation and infiltration of innate cells into the skin substantiate the use of the ATP alarmin model to assess the effects Trm cells have on innate infiltration and activation. To mirror *in vivo* conditions in which ATP release is continuous and to directly signal through the cutaneous P2X7R, BzATP (a specific P2X7R agonist and ATP analog) is co-administrated in the skin with an

ATPase inhibitor (POM1). This strategy directly targets P2X7R and prevents the endogenous breakdown of BzATP. Thus, utilizing the BzATP/P2X7R model, we compared the capacity of DM and SPF mice to induce an innate immune response in the skin. Consistent with our previous findings^21^, analysis by flow cytometry revealed a significant increase in neutrophils following treatment with BzATP + POM1 in SPF mice compared to PBS controls (**Figure 4A and 4B**). However, there was not a significant increase in neutrophils in DM compared to PBS controls (**Figure 4A and 4B**). Notably, there was a significant decrease in neutrophil infiltration in DM compared to SPF mice following P2X7R stimulation (**Figure 4A and 4B**). It should be noted that the neutrophil population increased in SPF mice is Ly6G^+^Ly6C^+^, which has been described as a mature neutrophil population that is not immuno-suppressive.^25,26^ Hence, utilizing two separate inflammatory models and different analytical methods, these data indicate that cutaneous Trm cells can suppress innate responses in DM following exposure to either a PAMP or a DAMP.

**Figure 4.**
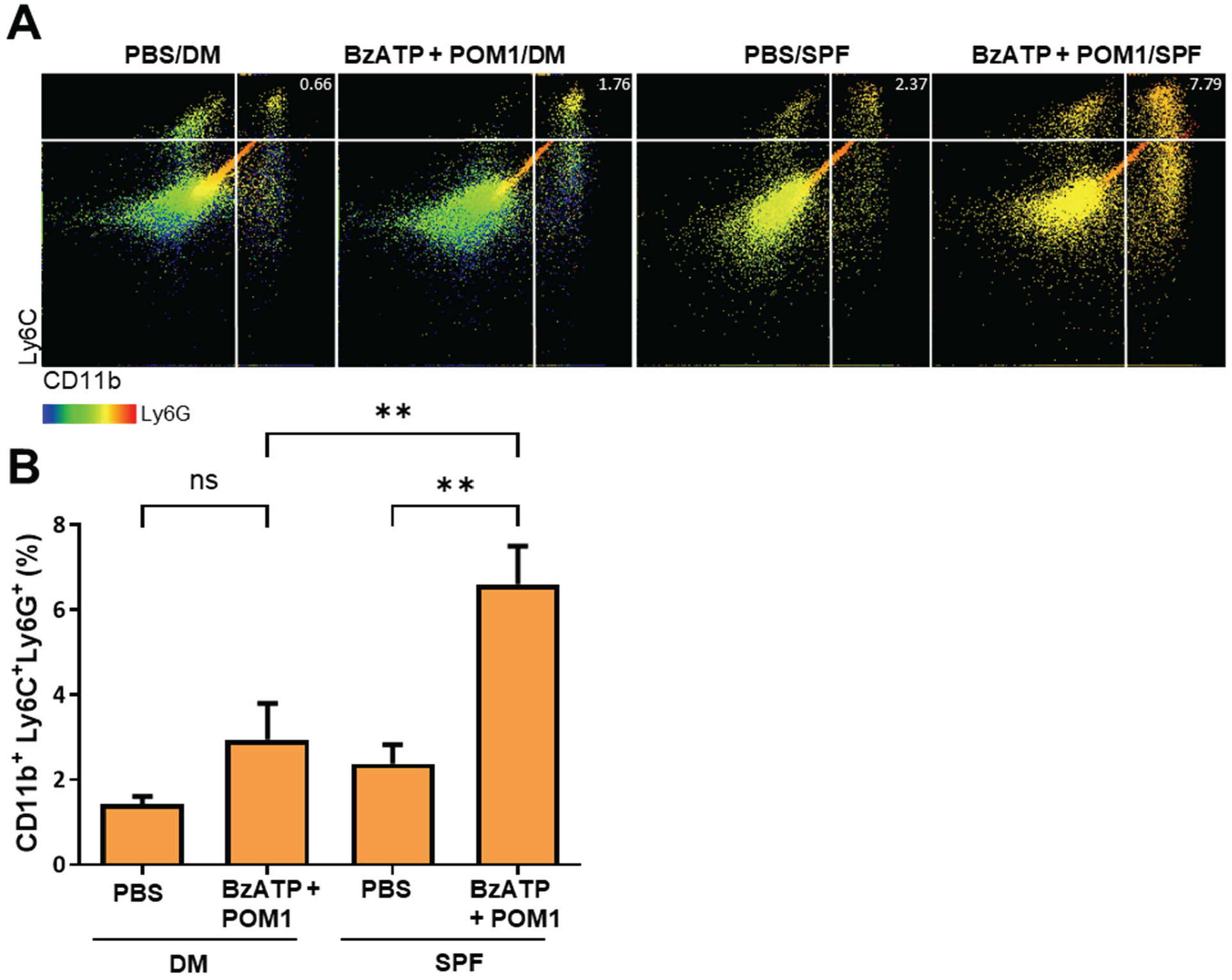
Neutrophils are decreased in DM following DAMP stimulation. Specific pathogen free mice (C57Bl/6 mice) or DM were injected with BzATP + POM1 (or PBS vehicle control) daily for 5 days then EOD for 4 days and sacrificed on day 10. (A) Phenotypic analysis of cutaneous infiltration was performed on day 10 by Aurora flow cytometry. Single-cell suspensions were assessed for LY6C+ and CD11b with 3^rd^ color analysis of Ly6G. Cells were initially gated on CD45^+^Via^-^, doublet exclusion, FSC vs SSC, and CD11c^-^. One representative of two independent experiments. (B) Flow cytometry data is quantitated. Bars are the mean ± SEM from 5-9 individual mice. Data are combined from 2 independent experiments.

IFN-γ has been demonstrated to block both neutrophil and monocyte migration.^12,13^ The mechanism of how IFN-γ blocks innate infiltration is not completely understood but is suggested to involve the control of chemokine expression.^12,13^ In this regard, we have determined that the T cell chemokine, Interferon gamma-induced protein 10 (IP-10/CXCL10) is significantly increased in DM compared to SPF mice following BzATP + POM1 treatments; conversely, the monocyte chemoattractant protein 1 (MCP-1/CCL2) is significantly decreased in DM compared to SPF mice (**Figure 5**). Thus, supporting the notion that cutaneous Trm cells have a role in orchestrating the immune response by attracting Tcm cells and suppressing innate cells. Likely, as an inherent mechanism to restrain aberrant inflammatory responses.

**Figure 5.**
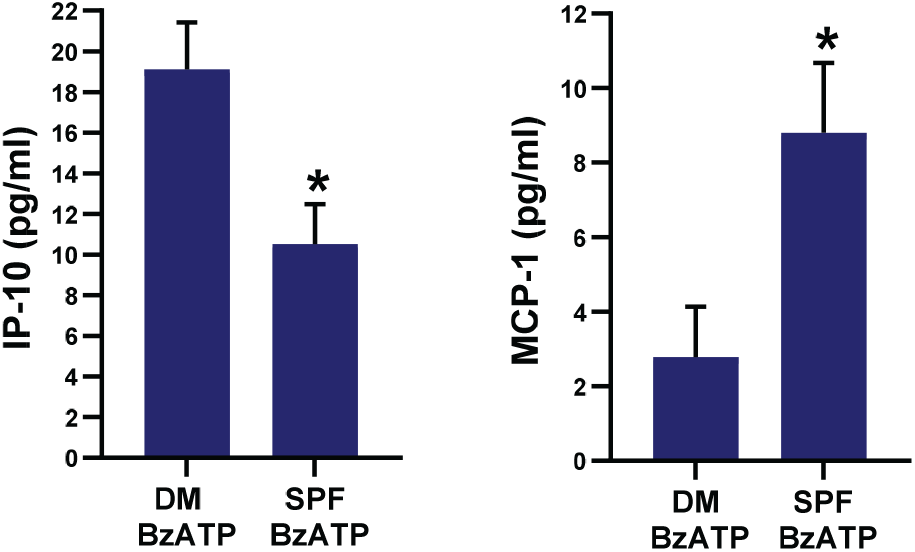
Interferon gamma-induced protein 10 is significantly increased in DM while MCP-1 is decreased. SPF and DM were injected with BzATP + POM1 (BzATP) daily for 5 days then EOD for 3 days and sacrificed on day 10. Serum was collected on day 10 and assessed by Luminex assays. Bars are the mean ± SEM from 8 individual mice. Each sample was run in duplicate. Data are combined from 2 independent experiments.

Psoriasis is a chronic inflammatory skin disease characterized clinically by red scaly plaques. The pathogenesis of psoriasis is largely dependent on IL-17 (IL-17A), which disrupts keratinocyte differentiation and indirectly induces neutrophil infiltration and neutrophilic microabscess formation.^27^ Trm cells capable of producing IL-17 reside in areas of resolved plaques and have been linked with the relapse of plaques in these sites.^1,28,29^ Given the above findings we set out to determine if non-lesional psoriatic skin populated with Trm cells would be capable of mounting an innate immune response following P2X7R stimulation. For these studies we utilized a xenotransplant model in which non-lesional biopsies were transplanted onto NSG mice. Two weeks following transplant grafts were injected with either PBS, as vehicle control, or BzATP + POM1. Interestingly, BzATP + POM1 could induce increased epidermal thickness, induction of neutrophilic microabscess formation, and cell clustering around the injection site (**Figure 6**). Indicating that in pathological conditions Trm cells have lost their capacity to suppress innate responses perhaps due to the complex interplay of IFN-γ and IL-17 responses. In this regard, CD49-CD8^+^ Trm cells secreting IL-17 in psoriatic skin have been described as having a prominent role in disease pathogenesis.^4,28^ Overall, the present studies have demonstrated that Trm cell populations can direct the outcome of the innate immune response depending on the microenvironmental cues.

**Figure 6.**
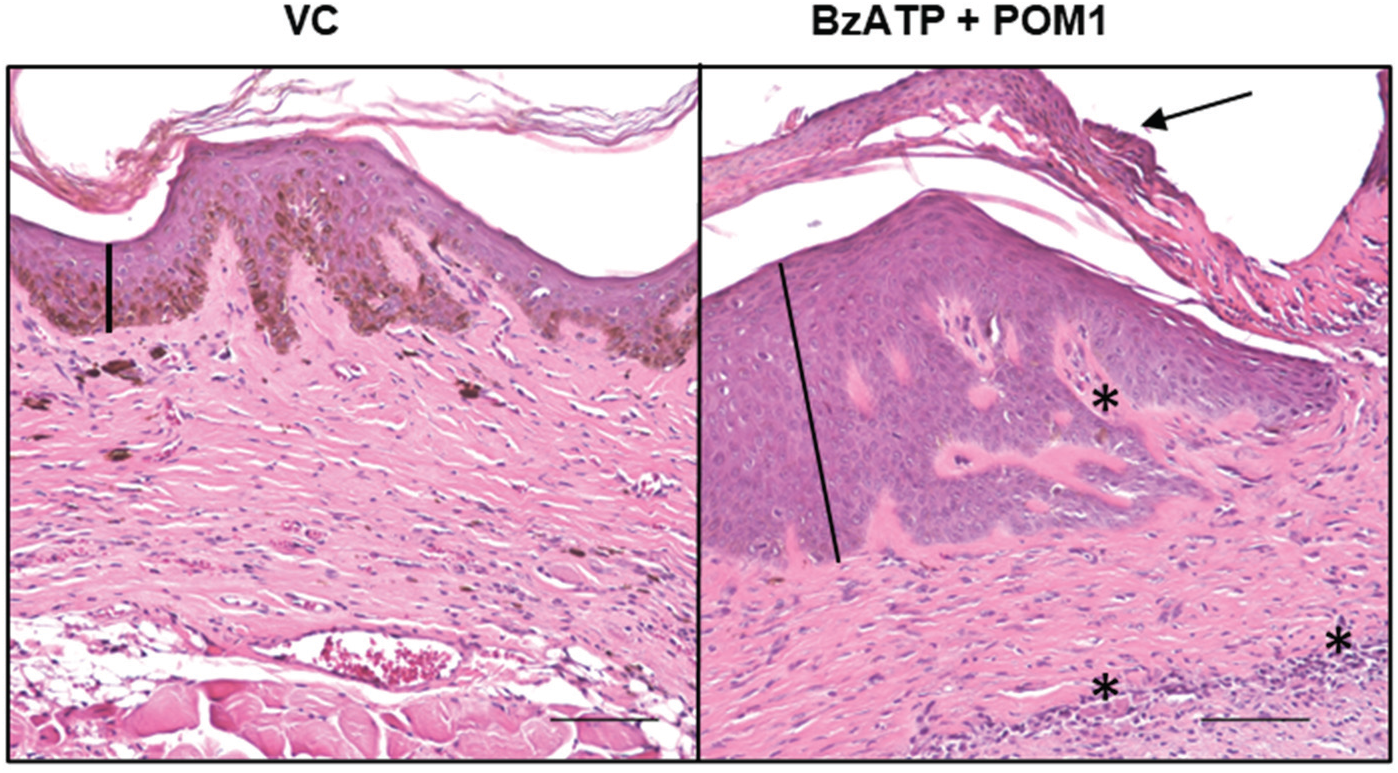
Psoriatic lesions develop in human skin xenotransplants following BzATP + POM1 injections. Two wks following xenotransplant of non-lesional human skin onto NSG mice, grafts were injected with BzATP + POM1 (or vanicream, **vehicle** control; VC) daily for 5 d. Two separate patients are represented. n=3 for each group. Black line indicates epidermal thickness, black arrow indicates neutrophilic microabscess, and asterisks represent cellular clustering.

## Acknowledgements

Research reported in this publication was supported by the National Institute of Arthritis and Musculoskeletal and Skin Diseases of the NIH under Award Numbers R01AR067746 (to ARM) and R21AR078349 (to ARM). The content of this manuscript is solely the responsibility of the authors and does not necessarily represent the official views of the National Institutes of Health.

